# *Irx3* Promotes Gap Junction Communication Between Uterine Stromal Cells to Regulate Vascularization during Embryo Implantation

**DOI:** 10.1101/2021.05.06.442331

**Authors:** Ryan M. Brown, Linda Wang, Anqi Fu, Athilakshmi Kannan, Michael Mussar, Indrani C. Bagchi, Joan S. Jorgensen

## Abstract

Spontaneous abortions have been reported to affect up to 43% of parous women, with over 20% occurring before pregnancy is clinically diagnosed. Establishment of pregnancy is critically dependent on proper embryo-uterine interactions at the time of implantation. Besides oocyte abnormalities, implantation failure is a major contributor to early pregnancy loss. Previously, we demonstrated that two members of the Iroquois homeobox transcription factor family, IRX3 and IRX5, exhibited distinct and dynamic expression profiles in the developing ovary to promote oocyte and follicle survival. Elimination of each gene independently caused subfertility, but with different breeding pattern outcomes. *Irx3* KO (*Irx3^LacZ/LacZ^*) females produced fewer pups throughout their reproductive lifespan which could only be partially explained by poor oocyte quality. Thus, we hypothesized that IRX3 is also expressed in the uterus where it acts to establish functional embryo-uterine interactions to support pregnancy. To test this hypothesis, we harvested pregnant uteri from control and *Irx3* KO females to evaluate IRX3 expression profiles and the integrity of embryo implantation sites. Our results indicate that IRX3 is expressed in the endometrial stromal cells of the pregnant uterus. Notably, of the days evaluated, IRX3 expression expanded into the endometrial stroma starting at day 4 of pregnancy (D4) with peak expression at D5-6, and then greatly diminished by D7. This pattern corresponds to the critical window for implantation and remodeling of the vasculature network in mice. Further, histology and immunohistochemistry at D7 showed that while embryos were able to attach to the uterus, implantation sites in *Irx3* KO pregnant mice exhibited impaired vascularization. In addition, our results showed significantly diminished expression of decidualization markers and disruptions in GJA1 organization in the decidual bed. These data, taken together with previous reports focused on the ovary, suggest that IRX3 promotes fertility via at least two different mechanisms: 1) promoting competent oocytes and 2) facilitating functional embryo-uterine interactions during implantation. Future research aims to tease apart the roles for IRX3 in the oocyte versus the uterus and the mechanisms by which it promotes early embryo survival and a successful pregnancy outcome.

## Introduction

Embryo implantation is achieved when a competent oocyte is fertilized and then develops into a blastocyst capable of facilitating embryo-uterine interactions with the receptive endometrium. In humans, it has been reported that approximately two thirds of pregnancies are lost due to implantation failure. Uncovering the signaling pathways and downstream mediators that govern implantation are necessary to improve outcomes associated with defective embryo implantation including ectopic pregnancies, implantation failure and infertility (reviewed in [1]).

In the mammalian uterus, steroid hormones estrogen and progesterone induce structural and functional changes during early pregnancy to support embryo implantation [2–5]. Of these changes, uterine stromal differentiation, or decidualization, is a critical response to embryo recognition that initiates extensive tissue remodeling for proper maternal-fetal interactions during early pregnancy. During decidualization, the endometrial stromal cells of the uterus undergo differentiation into decidual cells with unique biosynthetic and secretory profiles needed to promote tissue transformation and uterine vascular remodeling [6–8]. In the murine model, implantation begins on day (D)4 midnight, with the majority of decidualization occurring between D5-8, and establishment of pregnancy by D10-11 of gestation [9]. Complementary timing occurs in the human uterus where, for a short window of time, during the mid-secretory phase of each menstrual cycle, the uterus becomes “receptive” to embryo implantation and the decidualization process begins [10]. In both the mouse and human, decidualization promotes tissue remodeling and neovascularization, which are critical events for successful establishment of pregnancy. A major challenge is deciphering the signaling mechanisms governing successful maternal-fetal interactions during early pregnancy. To this end, it is necessary to identify and characterize factors that regulate decidualization and angiogenesis during embryo implantation.

The Iroquois factors are highly conserved proteins that have been implicated in patterning and embryogenesis in animal kingdoms spanning invertebrate to vertebrate. In mammals, there are a total of six Iroquois genes clustered in two groups, with cluster A (*Irx1,2,4*) and cluster B (*Irx3,5,6*) located on chromosomes 5 and 16 in the human and 8 and 13 in the mouse, respectively [11, 12]. Previously, we discovered that null mutation of both *Irx3* and *Irx5* resulted in abnormal granulosa cell morphology and disrupted granulosa cell-oocyte interactions [13]. A closer look, using *Irx3^LacZ/LacZ^* single knockout mice, revealed that although mutant females could become pregnant, loss of *Irx3* caused a decrease in birthrate to approximately half as many pups compared to controls throughout a 6-month breeding study. While *Irx3^LacZ/LacZ^* females demonstrated abnormalities in follicle survival, this did not fully explain the subfertility phenotype [13].

Although evidence points to an oocyte deficit with loss of *Irx3,* ovarian histology led us to hypothesize that another facet in the female reproductive axis could be disrupted. Due to the decrease in pup accumulation over time, we hypothesized that embryo implantation was compromised due to loss of *Irx3*. Evaluation of implantation sites within *Irx3^LacZ/LacZ^* pregnant females demonstrated impaired vascularization and a significant reduction in pups by D7 of pregnancy. Herein, we report for the first time that *Irx3* is expressed in the mouse uterus overlapping the window of implantation when it plays a critical role in neoangiogenesis. Our results suggest disruptions are caused by unorganized gap junction (GJA1) protein expression. Together, these findings reveal the multifaceted role of *Irx3* in female fertility.

## Materials and Methods

### Ethics Statement

Animals were euthanized by CO_2_ asphyxiation followed by cervical dislocation. Animal housing and all described procedures were reviewed and approved by the Institutional Animal Care and Use Committee at the University of Wisconsin – Madison and were performed in accordance with National Institute of Health Guiding Principles for the Care and Use of Laboratory Animals.

### Animals

Mice were housed in disposable, ventilated cages (Innovive, San Diego, CA). Rooms were maintained at 22 ± 2°C and 30–70% humidity on a 12-hour light/dark cycle. Mouse strains included CD1 outbred mice (Crl:CD1(ICR), Charles River, Wilmington, MA, USA) and *Irx3^LacZ^* [14], all of which were maintained on a CD1 genetic background. Genotyping was carried out as previously reported [13, 14]. Pregnancies were the result of breeding between CD1 (WT control) or *Irx3^LacZ/LacZ^* females and CD1 males. Thus, all embryos generated in *Irx3^LacZ/LacZ^* females were *Irx3^+/LacZ^*. Timed mating was identified as the presence of a vaginal plug after mating which was designated as day 1 (D1) of pregnancy.

### Tissue Processing and Histology

Uterine tissue was harvested at the indicated timepoints, fixed in 10% neutral buffer formalin (NBF) in phosphate buffer saline (PBS) at 4°C overnight. Uteri were collected on D7 of gestation, dehydrated through an ethanol gradient, cleared in xylene and embedded in paraffin. Paraffin blocks were sectioned at 5μm thickness, mounted on slides and then stained with hematoxylin and eosin (H&E) for histological analysis.

### Immunohistochemistry/Immunofluorescence

Uterus tissue sections were deparaffinized in xylene, rehydrated through a series of ethanol washes, and then rinsed in water. Antigen retrieval was performed by immersing the slides in 0.1M citrate buffer solution at pH 6.0 or EDTA buffer at pH 8 depending on antibody specifications (Table S1), and heated in a water bath at 80°C for 25 min. The slides were allowed to cool, rinsed in water, followed by PBS washes. Slides were then incubated at room temperature with 10% donkey solution for 1h before incubating them with primary antibody overnight at 4°C. The following morning, tissues were washed in PBS before incubating with secondary antibody for 1 hour, then washed with PBS, incubated with a 10X DAPI (4’,6-diamidino-2-phenylindole) in PBS solution (1:500) as a nuclear counterstain for 10 minutes, followed by 3 minutes with Vector True View Kit (Vector Laboratories, SP-8400, Burlingame, CA), and mounted. Immunofluorescence was repeated in uterine sections collected from at least 3 animals. Quantification of IHC was performed using Image J, the mean fluorescence intensity of CD31 (n=3 for each group) or GJA1 (n = 3-4 for each group) were quantified using the ROI manager on both the mesometrial and anti-mesometrial regions surrounding the embryo. Images were collected on a Leica SP8 confocal microscope for IHC and a Keyence BZ-X700 microscope for H&E.

### Quantitative real time PCR analysis (qPCR)

Uterine tissue was homogenized, and total RNA was extracted using TRIZOL reagent (Invitrogen, 15596026, Waltham, MA, USA), according to the manufacturer’s protocol and quantified using a NanoDrop 2000. 500ng from each sample was used for First-Strand cDNA synthesis by SuperScriptII (Invitrogen, AM9515, Waltham, MA, USA). cDNA was diluted 1:5 and then 2μl was added to 5ul SYBR green PCR mixture (BioRad, 1725271, Hercules, CA, USA), 2.4ul water and 1.25 pmol primer mix. PCR reactions were carried out using the BioRad CFX96 system and RNA transcripts were quantified using the ΔΔCt method (Livak & Schmittgen, 2001). Primers, *36b4* 5’ – CGACCTGGAAGTCCAACTAC – 3’ R: 5’ – ATCTGCTGCATCTGCTTG - 3’ and *Irx3* F: 5’ - CGCCTCAAGAAGGAGAACAAGA - 3’, R: 5’ - CGCTCGCTCCCATAAGCAT - 3’ (IDT, Coralville, IA, USA)

### Statistics

Statistics between groups were carried out using a two-sample t-test. Results were considered statistically significant if p-values were ≤ 0.05.

## Results

### *Irx3^LacZ/LacZ^* females exhibit defects in uterine vascularization with a marked reduction in implanted embryos

Previously, we determined that *Irx3^LacZ/LacZ^* females produced significantly fewer live pups over time compared to their controls, resulting in a subfertility phenotype. Ovarian histology and follicle counts identified oocyte deficits, but reproductive data suggested other factors were also contributing to the subfertility phenotype [13]. To understand the impact of *Irx3* on female fertility, we expanded our evaluation to examine uterine contributions of *Irx3* on fertility using *Irx3^LacZ/LacZ^* mice and their littermate controls over time. We investigated the integrity of implantation sites on D7 of pregnancy. Histological analysis of *Irx3^LacZ/LacZ^* (Fig 1B, D) and control (Fig 1A, C) implantation sites at D7 of gestation demonstrate that loss of *Irx3* impairs uterine vascularization, as depicted in the white boxes (Fig 1 A-D). To investigate whether impaired implantation and defective vascularity were a result of hormonal deficiency, we measured circulating estrogen and progesterone (Fig 1 E, F). Results showed that, estrogen and progesterone levels were comparable between *Irx3^LacZ/LacZ^* and litter mate controls. To assess when embryos were being lost in *Irx3^LacZ/LacZ^* females, we compared the number of implanted embryos to that of control mice on D7 and D11 of gestation (Fig 1 G). By D7 we saw a significant decrease in the number of pups *in utero* in the *Irx3^LacZ/LacZ^* mice; however, there was no further reduction in pups between D7 and D11 indicating that pups were being lost during the decidual phase of pregnancy. Together these data demonstrate that *Irx3* has a role in female fertility with implications in uterine angiogenesis during embryo implantation.

**Fig 1:**
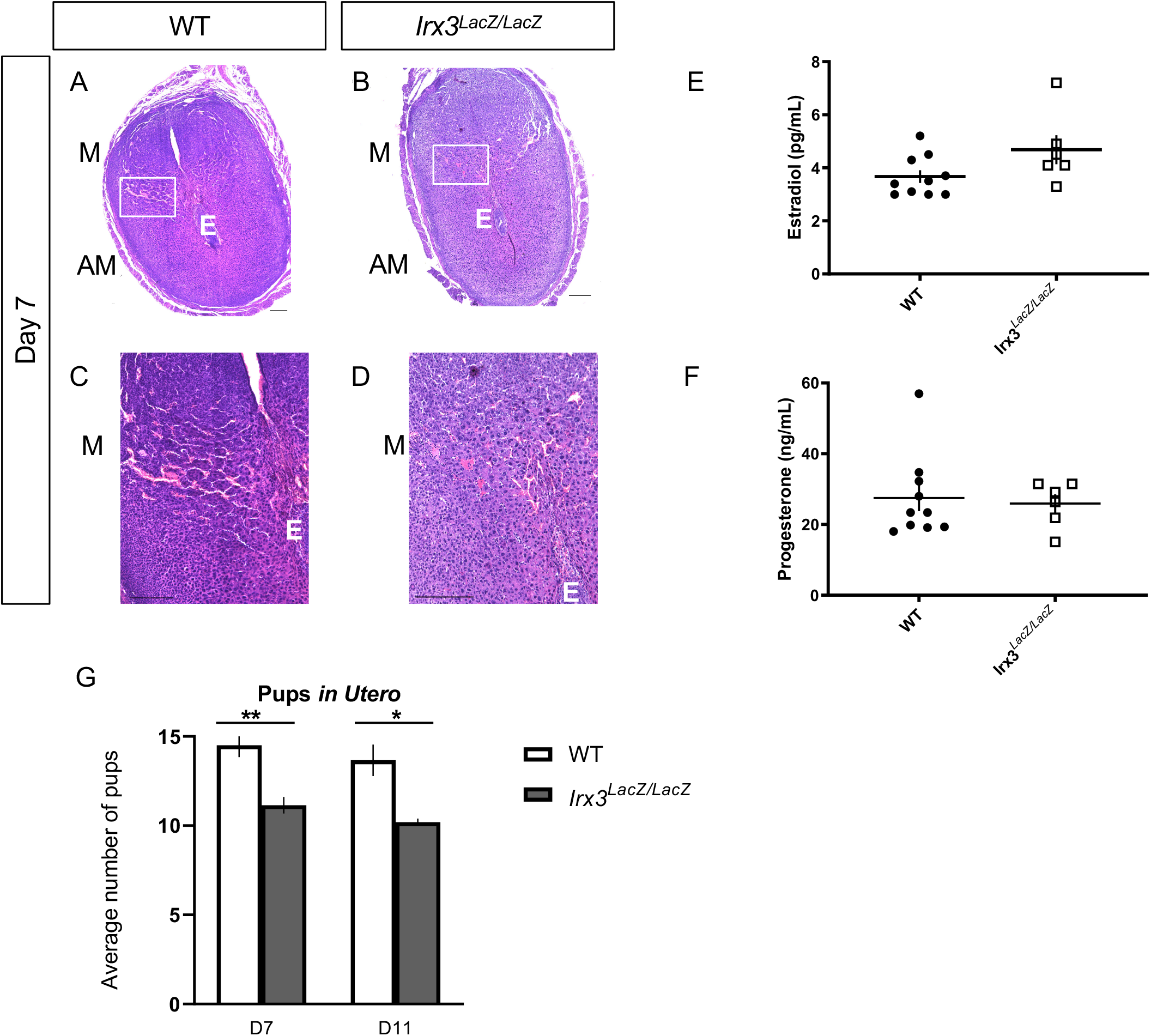
*Irx3^LacZ/LacZ^* causes deficits in uterine vascularization and subfertility. (A-D) H&E staining of D7 control (A, C n=6) and *Irx3^LacZ/LacZ^* (B, D n=6) pregnant uteri. Scale bars, 250μm. The white box in A and B are enlarged in C and D, respectively. A: anti-mesometrial; M: mesometrial; E: embryo. (E, F) Circulating Estrogen (E) and Progesterone (F) were measured at D7 of gestation (WT n=10; *Irx3^LacZ/LacZ^* n=6). (G) Average number of implantation sites *in utero* at D7 (WT n=8; *Irx3^LacZ/LacZ^* n=7) and D11 (WT n=3; *Irx3^LacZ/LacZ^* n=5). Data in G represents the mean ± SEM. Statistics: two-sample t-test, *: p<0.05; **: p<0.01.

### *Irx3* expression is confined to a discrete window during embryo implantation

Evidence of vascularity deficits within embryo implantation sites and early embryo loss suggested a role for *Irx3* in uterine implantation. Thus, we examined the protein and transcript profile of *Irx3* in the mouse uterus during normal pregnancy. Late on D4 of gestation, the mouse uterus is receptive, and implantation ensues. At this time IRX3 protein expression is detected predominately in the cytoplasm of the epithelial cells surrounding the lumen and uterine glands (Fig 2A, B). By D5, which is the onset of endometrial stromal cell decidualization, IRX3 protein expression is substantially increased, with expression expanding to the uterine stroma (S) immediately surrounding the embryo (E), also referred to as the primary decidual zone (Fig 2 C, D). Similarly, at D6 of gestation, IRX3 expression is prominently expressed throughout the decidualized stroma, expanding further into the secondary decidual zone (Fig 2 E, F). On D7 of gestation, nearing the end of the decidualization process, IRX3 protein expression is diminished (Fig 2 G, H). Analysis of *Irx3* transcripts demonstrate a complimentary profile with expression initially documented at D4, followed by substantial increases at the onset of decidualization at D5 and D6 and a sharp decline toward the end of decidualization on D7. These data demonstrate that *Irx3* is induced in endometrial stromal cells at the onset of embryo implantation, the expression peaks during decidualization, and declines with the cessation of decidual phase of pregnancy.

**Fig 2:**
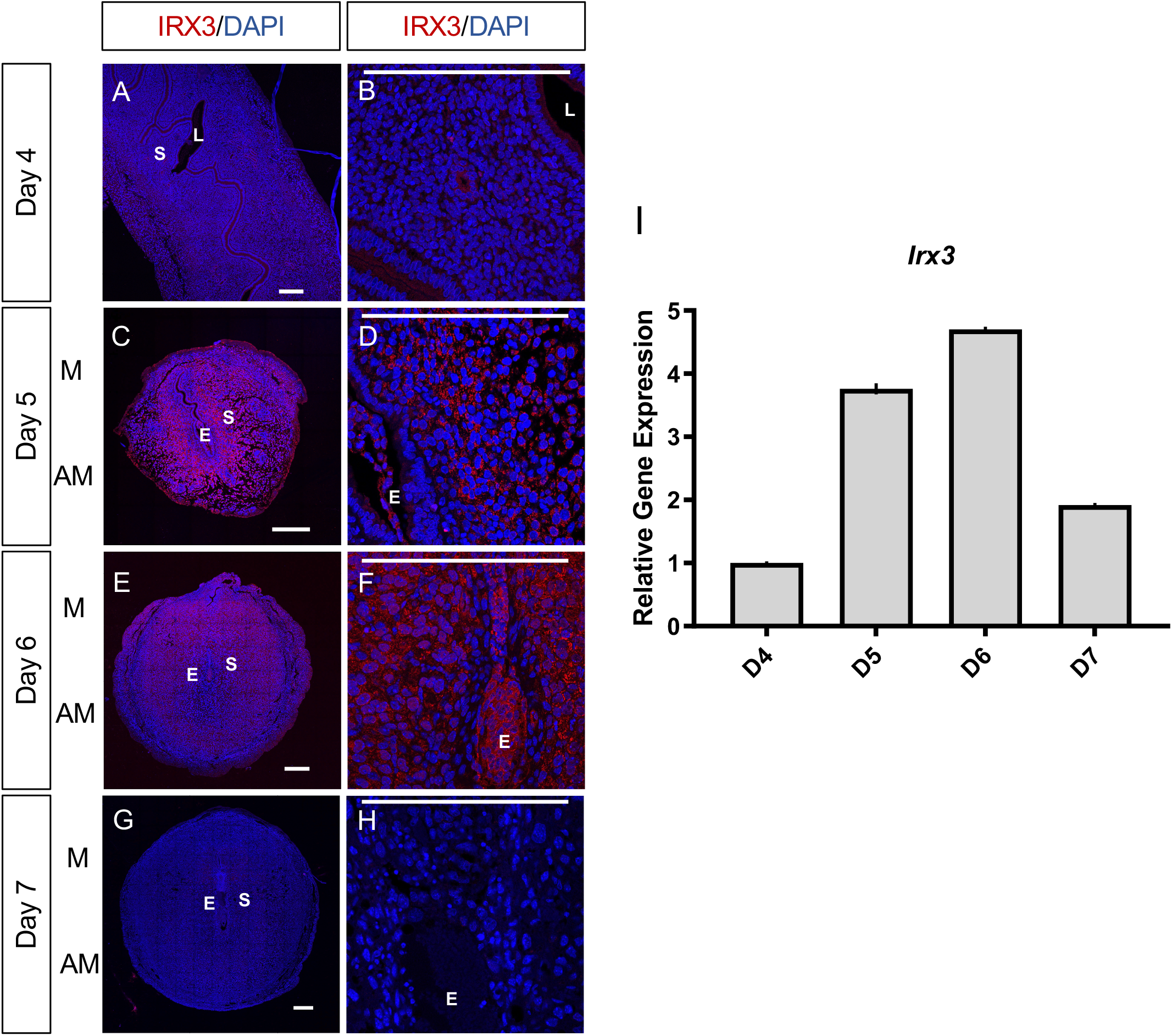
*Irx3* expression coincides with embryo implantation. (A-F) Immunofluorescence of IRX3 (red) co-labeled with DAPI, a nuclear marker (blue) throughout pregnancy days 4-7 in wild type mice implantation sites (n=3). Scale bars represent 250μm. L: lumen; S: stroma; E: embryo; M: mesometrial; A: anti-mesometrial. (I) Characterization of *Irx3* transcripts in the WT pregnant mouse uterus at days 4-7 (D4-7). Real time qPCR was determined by setting the expression level of *Irx3* mRNA on D4 of pregnancy to 1.0. Results are reported relative to 36b4 (n=3). Data represent the mean ± SEM of three biological replicates performed in triplicate at each time point.

### Endometrial stromal cell differentiation is impaired in *Irx3^LacZ/LacZ^* females

The close spatio-temporal relationship between *Irx3* expression and the progression of decidualization led us to hypothesize that *Irx3* may play a role in this process. We, therefore, evaluated the expression of known regulators of decidualization in *Irx3^LacZ/LacZ^* and control uteri. Implantation sites were harvested at D7 and processed for RNA isolation and immunofluorescence. Results show that as expected, *Irx3* expression was already low in WT uteri at D7 (set to 1) and detectable, but extremely low (75-80% decreased) in the mutant decidua. Any transcripts present were likely derived from embryo tissue contamination. While steroid receptor genes, *Esr1* and *Pgr1* remained unchanged, all other decidual markers were significantly decreased in mutant versus wild type implantation sites (Fig 3A). Further, HAND2 expression was weak in *Irx3^LacZ/LacZ^* D7 stroma compared to wild type as indicated by immunofluorescent analysis (Fig 3B), supporting qPCR results and suggesting defective decidualization.

**Fig 3:**
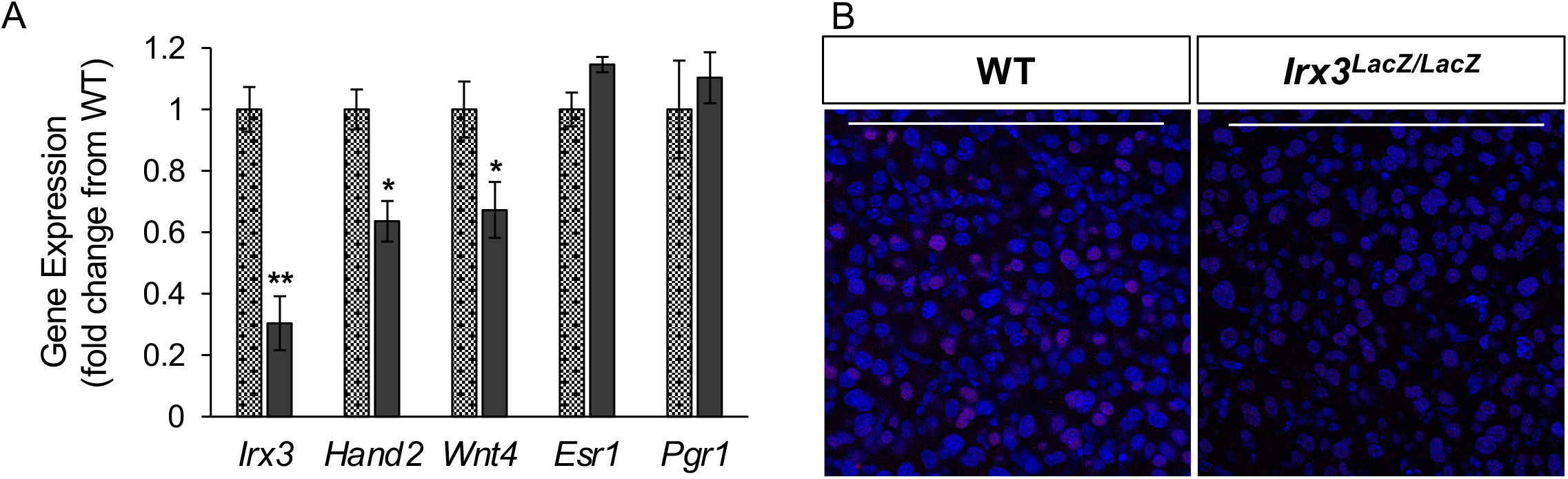
Decidualization is impaired in pregnant *Irx3^LacZ/LacZ^* uteri. (A) Real time qPCR was performed using total RNA isolated from implantation sites from pregnant uteri of WT (stippled light gray) and *Irx3^LacZ/LacZ^* (dark gray) females on day 7 of pregnancy. Data represents the mean ± SEM of three biological replicates performed in triplicate at each time point. Fold change was calculated relative to transcript levels of the WT sample. Statistics: Student’s t-test, *p ≤ 0.05. (B) Immunofluorescence of HAND2, a stromal cell marker (red) co-localized with DAPI, a nuclear marker (blue) at gestation D7 in control (n=4) and *Irx3^LacZ/LacZ^* (n=3) implantation sites. Scale bars represent 250μm.

### Compromised neoangiogenesis in the decidual bed of *Irx3* KO females

To further evaluate the vascularization defect found in *Irx3^LacZ/LacZ^* implantation sites, we examined whether loss of *Irx3* expression affected angiogenesis. Thus, we analyzed the angiogenic response in *Irx3^LacZ/LacZ^* uteri by monitoring an endothelial cell marker, CD31 (platelet endothelial cell adhesion molecule-1, PECAM-1), at D7 of gestation (Fig 4 A-G). As expected, implantation sites from control mice exhibited prominent CD31 expression in both the antimesometrial (AM) and mesometrial (M) areas on day 7 of pregnancy (Fig 4 A, C, E). In contrast, while expression of CD31 in *Irx3^LacZ/LacZ^* implantation sites was not different in the mesometrial side (Fig 4D, G), quantification of expression showed a significant decrease of CD31 expression in the antimesometrial region (Fig 4B, F, G). The decrease in the level of CD31 at the implantation site of *Irx3^LacZ/LacZ^* uteri on D7 raises the possibility that *Irx3* is mediating embryo-uterine interactions through promotion of a robust angiogenic response during implantation.

**Fig 4:**
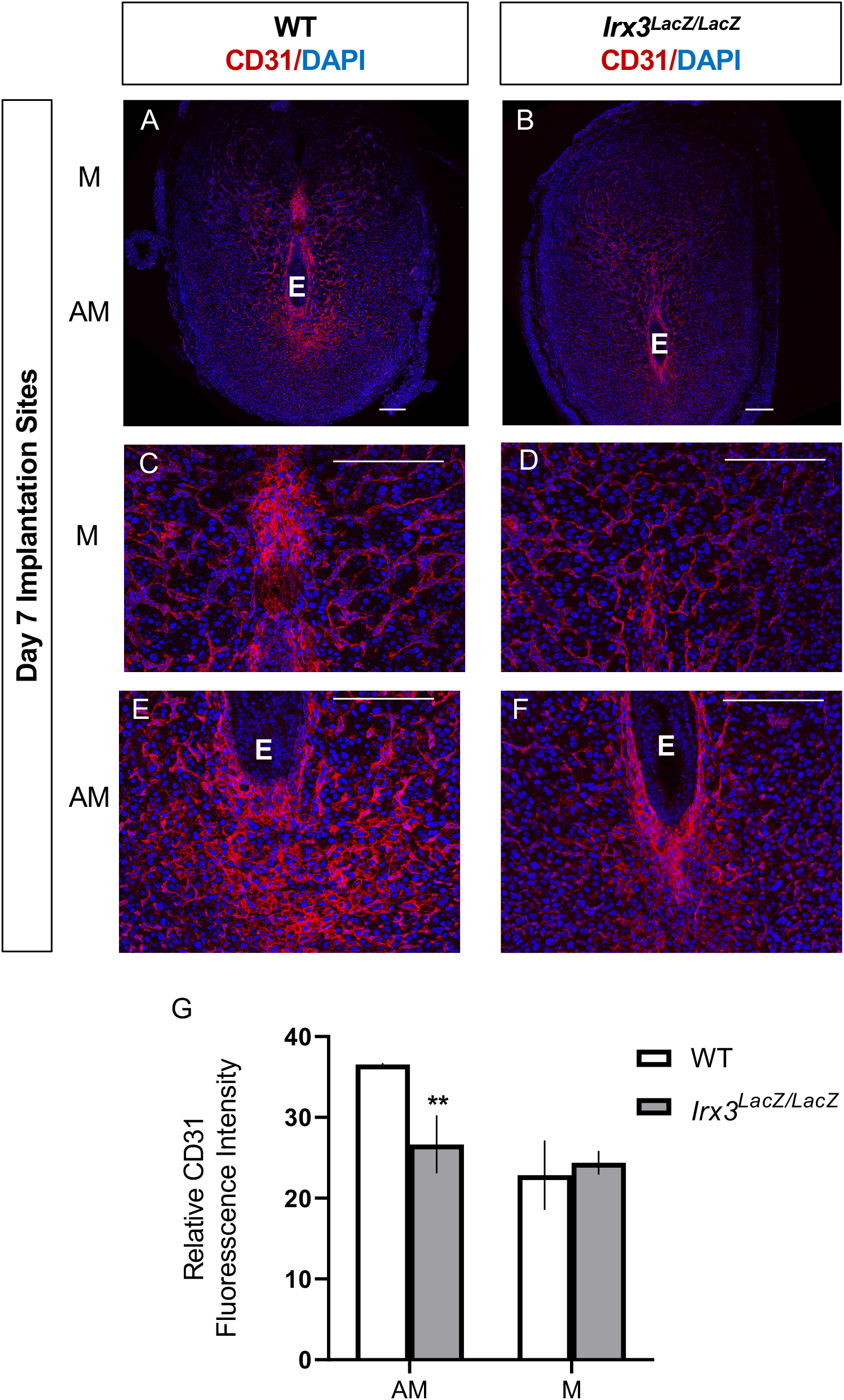
*Irx3^LacZ/LacZ^* uteri exhibit impaired vascularization by day 7 of gestation. (A-F) Immunofluorescence of platelet endothelial cell adhesion molecule (PECAM, CD31, red) co-labeled with DAPI, a nuclear marker (blue) at pregnancy D7 in control (A, C, E, n=3) and *Irx3^LacZ/LacZ^* (B, D, F, n=3) implantation sites. Scale bars represent 250μm. E: embryo; M: mesometrial; A: anti-mesometrial. (G) Quantification of relative CD31 fluorescence intensity in the anti-mesometrium (AM) and mesometrium (M). Data represents Mean ± SEM. Statistics: two-sample t-test; **: p<0.01.

### Fewer and unorganized gap junction connections in *Irx3^LacZ/LacZ^* uteri

To gain insights on the impact of *Irx3* loss on decidualization and vascularization, we investigated GJA1, a gap junction protein critical in modulating stromal differentiation and neovascularization during murine implantation [15]. In our previous investigations, *Irx3^LacZ/LacZ^* ovaries demonstrated no change in *Gja1* transcripts between mutant and control, but they detected abnormal deposition of GJA1, which resulted in follicle death [13]. Based on these data, we tested whether loss of *Irx3* in the uterus would impair GJA1 expression during embryo implantation. Complementary to our ovarian expression data, we found that loss of *Irx3* resulted in no change in Gja1 transcripts (Fig 5A), but significantly disrupted GJA1 protein localization in both mesometrial and anti-mesometrial regions of the uterine stroma (Fig 5 B-J). Together these data suggest that *Irx3* functions to promote embryo implantation through appropriate cell-cell communication to provide the proper environment for successful decidualization and neovascularization.

**Fig 5:**
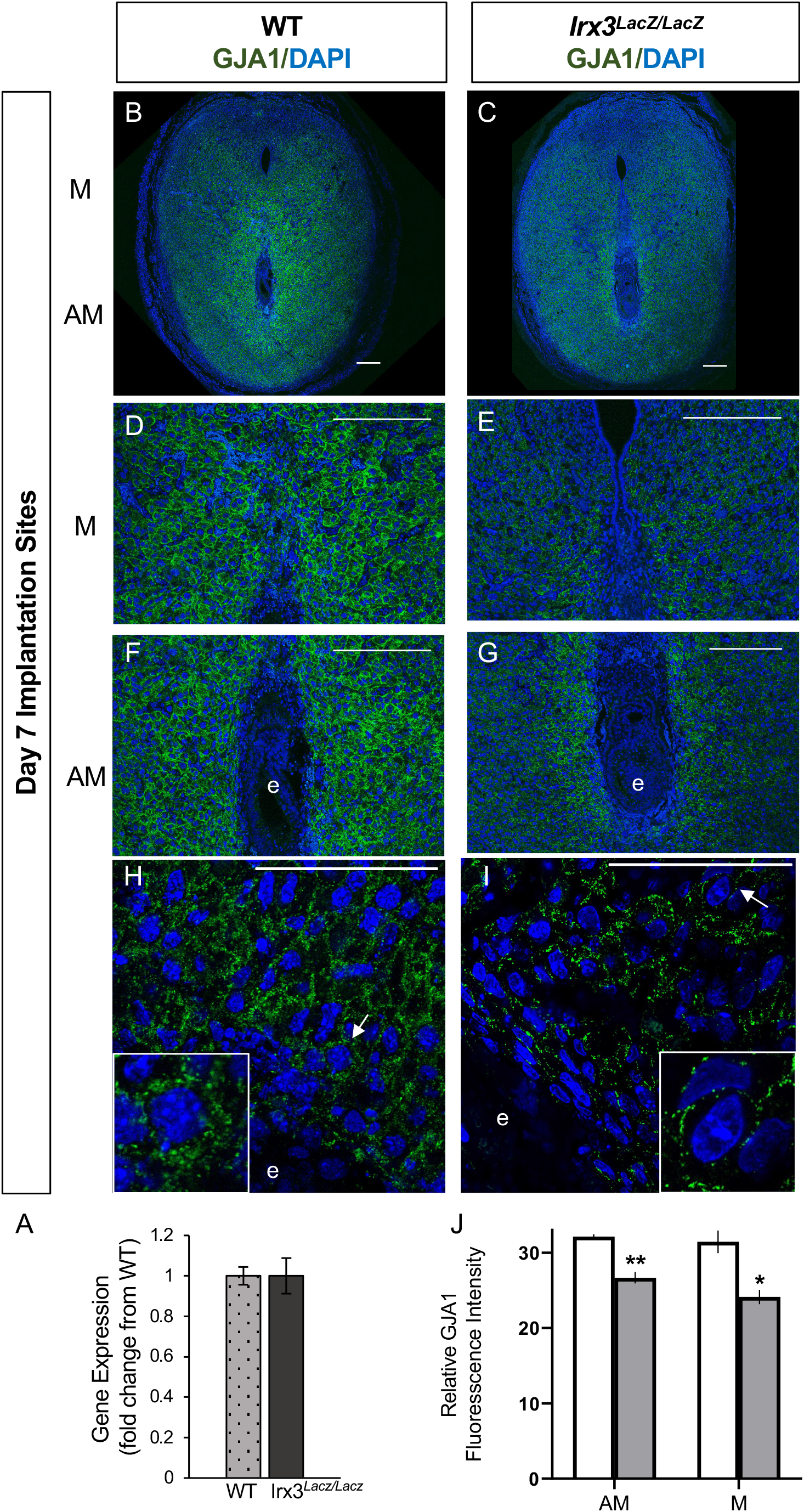
*Irx3^LacZ/LacZ^* uteri express similar *Gja1* transcripts but abnormal GJA1 protein. (A) Real time qPCR results for *Gja1* from pregnant uteri of WT and *Irx3^LacZ/LacZ^* females on D7 of pregnancy. Data represents the mean ± SEM of three biological replicates performed in triplicate at each time point. Fold change was calculated relative to transcript levels of the WT sample. Statistics: Student’s t-test, *p ≤ 0.05. (B-I) Immunofluorescence of gap junction protein 1 (connexin 43, GJA1, green) co-labeled with DAPI, a nuclear marker (blue) at gestation D7 in control (B, D, F, H, n=4) and *Irx3^LacZ/LacZ^* (C, E, G, I, n=5) implantation sites. Images are represented in increasing magnification, scale bars represent 250mm. Arrows highlight single cells that are enlarged within the inset of H and I. (J) Quantification of relative GJA1 fluorescence intensity in the anti-mesometrium (AM, p = 0.01) and mesometrium (M, p = 0.027) in wild type (white bars) vs *Irx3^LacZ/LacZ^* females (light gray bars). Data represents Mean ± SEM. Statistics: two-sample t-test, *: p<0.05; **: p<0.01.

## Discussion

The Iroquois genes have been implicated in embryonic patterning and can be found in a range of tissues, with roles in organization of the spinal cord, limb, bone and heart, to name a few [12, 16–18]. Previously, we identified IRX3 and IRX5 expression in mouse ovaries, but not testis, starting after sex determination [19]. This led to a series of investigations to discern the role of IRX3 and IRX5 in the ovary during development. Using mutant mice lacking both *Irx3* and *Irx5*, we discovered mutant ovaries had abnormal granulosa cell morphology and disturbed granulosa cell-oocyte interactions [13]. A closer look at ablation of *Irx3* alone, using *Irx3^LacZ/LacZ^* mutant females, indicated that loss of *Irx3* caused reduced follicle numbers; however, this did not fully explain the profound deficit in fertility as indicated by at least 50% loss in live pup births compared to wild type females [20]. This prompted us to investigate whether IRX3 was affecting embryo implantation. Results from the current study suggest multifaceted roles for IRX3 in female fertility that include important functions in both ovary and pregnant uterus. In particular, we report that IRX3 functions in early pregnancy to establish successful embryo-uterine interactions. These data provide a foundation for new discoveries of how IRX3 functions in a spatiotemporal manner in the uterus to promote successful embryo-uterine interactions via establishment of proper cell-cell communications to support both decidualization and angiogenesis.

Establishment of a robust and healthy vascular network is essential for reproduction as it supports folliculogenesis, ovarian hormone production, ovulation, implantation and embryonic growth [21–23]. Evaluation of the integrity of *Irx3^LacZ/LacZ^* implantation sites on D7 revealed reduced vascularity with a concomitant decline in the number of implanted embryos, indicating disruptions in reproductive fecundity in *Irx3^LacZ/LacZ^* mice. Together these data demonstrate that *Irx3* has a role in uterine angiogenesis during embryo implantation.

In the murine model, implantation begins late on day D4 when the mouse uterus is receptive to implantation [24]. At D5, the decidualization process begins and the stromal cells of the uterus differentiate into a unique secretory tissue, the decidua. The stromal cells immediately surrounding the embryo are transformed into the primary decidual cells and further expand into the secondary decidual cells until the invasive period is complete [9]. Here, we demonstrate that the expression profile of IRX3 in uterine stromal cells is intimately associated with the decidualization phase of mouse pregnancy. Notably, during the mid-secretory phase of the human menstrual cycle, studies show that *IRX3* expression increases as the uterus becomes receptive to implantation [25–27]. Contrary to mice, the human uterus begins decidualizing in the nonpregnant state, during the mid-secretory phase with an expansion of decidualization once the pregnancy is established [28]. Taken together, these data and our studies have uncovered a conserved physiological timing of *Irx3* expression in the human and murine uterus that corresponds to the onset of decidualization in both species.

As decidualization progresses, the uterus undergoes expansive tissue remodeling, critical for proper maternal-fetal interactions. Following the attachment of the blastocyst to the uterine epithelium, the underlying stromal cells differentiate into decidual cells with unique biosynthetic and secretory profiles needed to promote this transformation. We found that IRX3 is robustly induced in the decidual tissue during a critical time frame in mouse implantation and angiogenesis. Factors secreted by the decidualizing stromal cells promote tissue and vascular remodeling, thus, it is conceivable that IRX3, produced by the decidualizing stromal cells, has a role in preparing the uterus for appropriate embryo-uterine interactions. Examination of the decidual program in *Irx3^LacZ/LacZ^* uteri indicated that initial aspects of the decidualization process are impaired in these mice. It has been suggested that stromal differentiation and angiogenesis are intimately connected processes during pregnancy [15]. Along with a difference in the presence of a subset of decidualization markers, we also observed a substantial decrease in CD31 expression, an endothelial cell marker, in the *Irx3^LacZ/LacZ^* implantation sites. Collectively, the reduction in vascularization seen at D7 by both histological analysis and immunohistochemistry, demonstrates that ablation of *Irx3* leads to improper vascularization in the mouse uterus during early pregnancy.

Previously, we identified that loss of both *Irx3* and *Irx5* impaired gap junction protein deposition in ovarian follicles, leading to follicle and oocyte death [13]. Here, we discovered a potentially conserved role for *Irx3* in mediating proper cell-cell communication via gap junction expression during embryo implantation. Early studies identified 3 connexins in the human uterus; connexin 26, connexin 32, and connexin 43 [29]. Of these, connexin 43 (GJA1) is the predominant subtype expressed in human and mouse endometrium and localized almost exclusively to the stroma, connecting decidual cells [30]. Conditional knockout of *Gja1* in the mouse uterus resulted in severe subfertility with aberrant differentiation of the uterine stroma and significant reduction in VEGF [15]. Further, maternal decidua isolated from women with recurrent early pregnancy loss was found to have reduced *Gja1* transcripts and protein when compared to controls [31]. The importance of GJA1 regulation during pregnancy is augmented by epidemiological studies that identified taking mefloquine, an anti-malarial compound that blocks GJA1, as a risk factor for spontaneous abortion [32]. Here, our data indicates that IRX3 is instrumental in mediating proper cell-cell communication via GJA1 expression. In the uterus and ovary, GJA1 serves a critical role in establishing communications between cells, promoting both oocyte competence in the ovary and angiogenesis in the uterus during pregnancy [15, 33, 34]. Our results indicate that IRX3 promotes successful cell-cell connections allowing for appropriate intercellular cross talk throughout the intimate correspondence of embryo implantation and angiogenesis during pregnancy.

Embryo implantation is achieved when a competent oocyte develops into a blastocyst capable of facilitating embryo-uterine interactions with the receptive endometrium. It is clear that IRX3 has a significant role in mouse reproductive health including oocyte competence and endometrial angiogenesis [15, 33, 34]. Impaired uterine receptivity and poor vascularization are major causes of early pregnancy loss. Therefore, understanding how *Irx3* is regulated during this critical window of implantation may provide insights and solutions for female reproductive health and fertility.

## Acknowledgements

We thank all members of the Jorgensen laboratory for comments and support. We thank Emily Huber and Eowyn Liu for their help with sectioning. We truly appreciate all of the comments, advice, and technical support from the Developmental Endocrinology Group at the University of Wisconsin – Madison School of Veterinary Medicine. We are immensely grateful for Miranda Sun and Daniel Radecki for their support in microscopy trainings and Image J. We would also like to show our gratitude to Annie Novak and Tori Gronemeyer for their exemplary mouse husbandry work.

## Supplementary Table

**Supplementary Table S1:** Antibody Information

| Antibody | Dilution | Company | Catalogue number | Antigen Retrival |
| --- | --- | --- | --- | --- |
| IRX3 | 1:100 | Invitrogen | PA5-35149 | EDTA |
| HAND2 | 1:50 | Abcam | ab200040 | Citrate |
| FRA1 | 1:100 | Invitrogen | PA5-76185 | Citrate |
| CD31 | 1:200 | Abcam | ab182981 | EDTA |
| GJA1 | 1:250 | Abcam | ab11370 | Citrate |

## Notes

**Funding** National Institutes of Health, Eunice Kennedy Shriver National Institute Of Child Health & Human Development (NIH-NICHD) R01HD075079 (JSJ) National Institutes of Health, Eunice Kennedy Shriver National Institute Of Child Health & Human Development (NIH-NICHD) R01HD090066 (IB) Endocrinology and Reproductive Physiology Training Grant, and National Institutes of Health (NIH-NICHD) T32HD41921 (AF).

### Competing Interest Statement

The authors have declared no competing interest.

